# Management of a Walnut Germplasm Collection: Which of SSR or SNP Markers Are Most Suitable to Preserve Biodiversity?

**DOI:** 10.1101/2019.12.17.879627

**Authors:** Anthony Bernard, Teresa Barreneche, Armel Donkpegan, Lheureux Fabrice, Elisabeth Dirlewanger

**Author notes:** Corresponding author: Elisabeth Dirlewanger.

## Abstract

The preservation of the maximum of diversity within the smallest number of accessions is one of the challenges of germplasm management. To construct core-collections, the assessment of the population structure and the relationships between the accessions represents a key step and the choice of suitable molecular markers is the starting point. Since the expansion of available SNP-based genomics tools, a debate has emerged regarding the usefulness of the widely used microsatellites (SSRs) markers. In this study, we analysed a part of the INRAE walnut germplasm collection of 150 accessions, unique in Europe for walnut biodiversity conservation, by comparing the power of both types of marker. We found that the first level of structure is equally detected using 13 SSRs or the Axiom™ *J. regia* 700K SNP array, and is in relation with the geographical origin of the accessions. For K=2, there was no exchange of accession between the two groups when both markers were compared. We also highlighted empirically that approximately 100 SNPs are needed to obtain similar clustering to SSRs in Principal Coordinate Analysis (PCoA). The neighbor-joining trees constructed were also consistent between both types of marker. The main differences lied in the upper levels of structure from K=3 to K=6, more powerful using the SNPs, and in the percentage of the explained variation in PCoA for K=2, higher using SSRs. We then constructed core-collections of 50 accessions, a crucial step in genetic resources management to reduce the costs and preserve the allelic diversity. Using two different construction methods, both SSR and SNP markers were suitable and able to keep at least 88.57% of the alleles. 32/50 accessions were in common between the two markers, for both methods. We concluded that the use of either marker is dependent on the researcher’s goal.

## Introduction

In the context of climate change and human population growth, plant genetic resources (PGR) are of upmost importance and they are facing crucial issues, since they constitute the foundations of the agricultural sustainability and the global food safety [1, 2]. Over the last three decades, the increase in the number of studies carried out on the discovery of new PGR and the exploration of existing ones has led to the production of numerous phenotypic and genotypic “big data” that are assessed to increase the effectiveness of their conservation and use [3]. This has raised questions about the governance systems of these resources and the exchange of materials, and therefore has led to the International Treaty on Plant Genetic Resources for Food and Agriculture (ITPGRFA), promoting the introduction of Digital Object Identifier (DOI) for each PGR, and standardization protocols for their characterization. It has been followed by the Nagoya Protocol, adopted in 2010, on the “access to PGR and the fair and equitable sharing of benefits from their utilization”.

The awareness of the need to share and characterize PGR does not solve the problem of genetic erosion. According to the Food and Agriculture Organization of the United Nations (FAO) 2010 Second Report on the state of the world’s PGR for food and agriculture, they are an estimated 7.4 million accessions (more than 28% of which are wheat, rice and barley) held in 1,625 banks. Nevertheless, FAO highlights a mixed picture. For example, the number and coverage of protected areas has increased by 30% over the past decade, increasing the level of protection of wild species of cultivated plants, but progresses are still needed outside these areas. Regarding *ex situ* management of PGR, they are mainly as seeds and some collections are at risk, due in part to the fact that they are generally underfunded, and that evaluation and characterization are often imprecise or inadequate [4]. In that respect, PGR management carried out with care is crucial from storage to use [5]. For clonally propagated perennial species, the conservation of PGR is generally done in *ex situ* orchards as grafted cultivars, which has pros and cons: the main advantages are that they can be stored under the climate conditions of their intended use, and can be evaluated during storage; but on the other hand, they require a lot of space, the cost of conservation is significant [6].

Nowadays, molecular tools contribute to each step of PGR management [7], since they can assist to find genetically close or synonym accessions to create “core collection” which will contain the maximum of genetic diversity within the smallest number of accessions, leading in particular to the reduction in conservation costs. They can be used then to decipher the genetic bases of agronomic traits, and used in selection processes. Before the development of genomics tools, now based mainly on biallelic Single-Nucleotide Polymorphisms (SNPs), the frequently used multiallelic Simple Sequence Repeats (SSRs) had become the markers of choice because of their high polymorphism. In Persian walnut (*Juglans regia* L.), a widely disseminated and grown species in many temperate regions, more than 20 publications mention the use of SSRs [8]. Recently, a high-density Axiom™ *J. regia* 700K SNP genotyping array was developed and validated, initiating a novel genomic area in walnut [9].

As a result, a legitimate debate arised about the consistency of the results found, and the type of marker that should be preferentially used to conduct population structure analysis or tasks related to germplasm management. Neutral SSR loci, due to slippage during replication, usually mutate much more frequently than SNP loci, leading to population-specific alleles useful to reveal population structure [10], but they therefore could not reflect the genome-wide genetic diversity [11]. In contrast, SNP loci are much more frequent in the genome of most species. These two types of marker can bring different views of the structure, and the merits of each are listed in [12]. Some other works focused on SSR and SNP comparisons in short-lived species such as rice [13], maize [14–16], sunflower [17], bean [18], and cowpea [19], to assess population structure and relatedness. Examples are rarer in perennials but exist in grape [20], and jujube [21]. By comparing those works, it is noticed that results found are conflicting. Moreover, if example of construction of core collections using both SSRs and SNPs is reported [22, 23], knowledge is still lacking regarding their comparison for this specific purpose.

Based on the walnut germplasm collection of the Institut National de Recherche pour l’Agriculture, l’Alimentation et l’Environnement (INRAE), the aim of this paper is to compare (*i*) the structure and relatedness among accessions using SSR or SNP markers, (*ii*) the core collections based on SSR or SNP markers using two building methods.

## Materials and Methods

### Plant materials and DNA extraction

The panel of the study consists of 150 unique accessions of *Juglans regia* from worldwide maintained at the *Prunus* and *Juglans* Genetic Resources Center, and located in the Fruit Experimental Unit of INRAE in Toulenne, France (latitude 44°34’37.442”N – longitude 0°16’51.48”W), near Bordeaux (Table S1). The INRAE walnut germplasm collection is a result of important collecting work performed between 1988 and 2000 in 23 countries including the European, American, and Asian continents.

The panel choice was made thanks to a previous work based on genetic diversity results using SSRs, and phenotypic variability [24]. The genomic DNAs of the panel were extracted from young leaves as described in this previous work.

### Genotyping using SSR and SNP markers

The panel was genotyped using 13 neutral SSR markers as described previously [24] and 609,658 SNPs from the Axiom™ *J. regia* 700K SNP array uniformly distributed over the 16 *J. regia* chromosomes [9]. The quality control steps were performed using “PLINK 1.9” software [25]. Poly High Resolution (PHR) and No Minor Homozygotes (NMH) SNPs were filtered using stringent thresholds: SNP call rate (> 90%), minor allele frequency (MAF > 5%), and redundancy in the genome (SNP probes aligning in duplicated regions). Finally, 364,275 robust SNPs (59.8% of the total number of SNPs) were retained for the following analyzes.

### Structure analyzes and core collection construction

Principal Coordinate Analysis (PCoA was used to determine the patterns of structure among the 150 accessions. Dissimilarities, based on allelic data, were calculated with 10,000 bootstraps, and transformed into Euclidean distances using a power transformation of 0.5. PCoA was performed using “DARwin 6.0.14” software [26], supplemented by “scatterplot3d” R package for 3D visualization.

As linked SNPs can account for too much in the population structure variance, particularly in linkage disequilibrium (LD) regions [27], a pruned subset of SNPs was also used for the PCoA. This filtering was completed using “PLINK 1.9” software, keeping only the SNP with the higher minor allele frequency, and based on a threshold of r²=0.2 (command --indep-pairwise 50 5 0.2). A subset of 100 SNPs, number similar to the number of SSR alleles, randomly selected, was also tested using “PLINK 1.9” software (command --thin-count 100) to compare PCoA results.

Genetic structure of our panel was also investigated using two types of analyses depending on the markers. Bayesian model-based analysis using “STRUCTURE 2.3.4” software was implemented for the SSR markers [28]. To identify the best number of clusters (K), ten runs were performed by setting K from 1 to 7. Each run consisted of a length of burn-in period of 100,000 followed by 750,000 Markov Chain Monte Carlo (MCMC) replicates, assuming an admixture model and correlated allele frequencies. The ΔK method [29], implemented in “STRUCTURE harvester” [30] was used to determine the most likely K.

Using SNP markers, sparse non-negative matrix factorization algorithm was implemented using “sNMF 2.0” software, available as a function of the “LEA” R package [31]. This software presents a fast and efficient program for estimating individual admixture coefficients from large genomic data sets, and produces results very close to Bayesian clustering programs such as “STRUCTURE” [32]. The choice of the best number of clusters (K) is based on a cross-entropy criterion implemented in “LEA” R package. For SSR and SNP markers, thresholds of 0.8 and 0.7 for admixture coefficient, respectively, were chosen to consider one accession as admixed.

Then, the genetic relationships between the 150 accessions were also assessed by the Neighbor-joining method [33] using “DARwin 6.0.14” software. The Unweighted Neighbor-Joining option was used to build the trees. In addition, core collections were constructed with a sampling intensity of 33% (n=50/150), using two methods: (*i*) the “maximum length sub tree” function [34] of “DARwin 6.0.14” software, which looks for a subset of accessions minimizing the redundancy between them, and limiting if possible the loss of diversity (the diversity here is expressed by the tree built), and (*ii*) the “entry-to-nearest-entry” method [35], implemented in “Core Hunter 3” software [36], which looks for a subset of accessions as different as possible from each other, avoiding selecting a few clusters of similar accessions at the extreme ends of the distribution. The number of alleles retained in one core collection was estimated.

## Results

### First level of structure analyzes

The most likely K subpopulations were evaluated considering the ΔK method and the cross-entropy criterion by using SSR and SNP markers, respectively. Using SSR markers, the higher drop of ΔK is for K=2, then followed by a raise for K=6 (Figure 1a). Very close findings are found using SNP markers, since the higher drop of the cross-entropy criterion is for K=2, with a curve slope starting for K=6 (Figure 1b).

**Fig 1.**
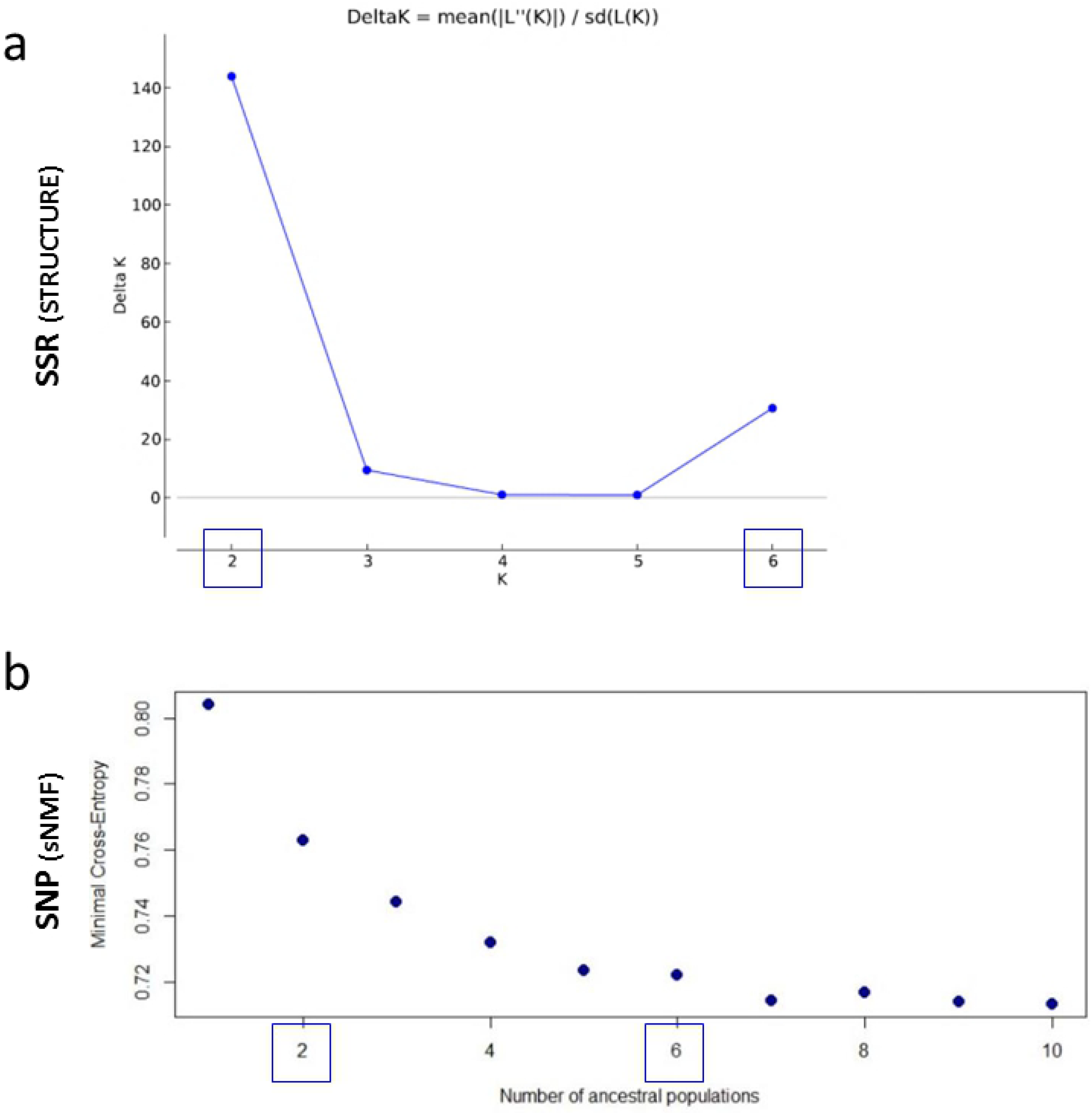
The most likely K subpopulations. K was evaluated considering a) the ΔK method by using SSR, and b) the cross-entropy criterion using SNP markers.

We assessed the first level of structure for K=2 to compare the results using SSR and SNP markers (Table S2). The individual admixture coefficients are showed for each of the 150 accessions (Figure 2). For both marker types, the clustering is linked to the geographic origin of the accessions. The group A contains the accessions from “Eastern Europe and Asia” (named “E”), from Afghanistan, Bulgaria, China, Greece, India, Iran, Israel, Japan, Romania, Russia, and Central Asia. The group B contains the accessions from “Western Europe and America” (named “W”) from Austria, Chile, France, Germany, Hungary, Netherlands, Poland, Portugal, Serbia, Slovenia, Spain, Switzerland, and USA and hybrids from INRAE.

**Fig 2.**
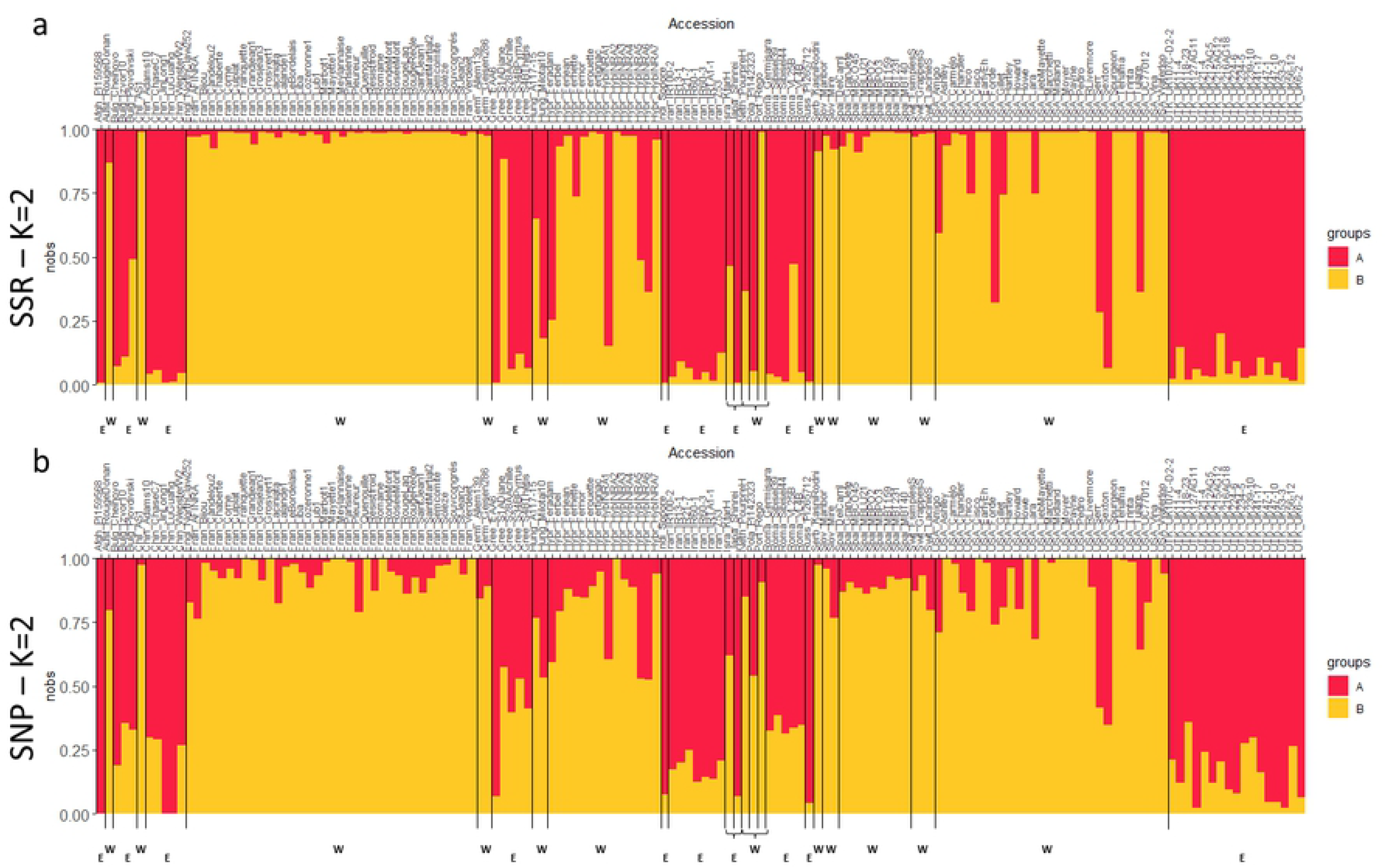
The bar plots showing the individual admixture coefficients of the 150 accessions for K=2. Structure was assessed a) using SSR, and b) using SNP markers. The accessions are ordered by their country of origin, by alphabetical order. The group A in red contains the accessions from Eastern Europe and Asia (“E”), whereas the group B in yellow contains the accessions from Western Europe and America (“W”) and hybrids from INRAE.

Using SSR markers, with a threshold of 0.8 for individual admixture coefficient, we found 17 admixed accessions, so 88.7% of the accessions were assigned to a group (Table 1). They are mainly French (‘Feradam’, ‘Fernette’, ‘Hybrid INRA 5’, ‘Hybrid INRA 6’), and California (‘Lara’, ‘Serr’, ‘Chico’, ‘Amigo’, ‘Gillet’, ‘Forde’, ‘Tulare’) modern hybrids. We also found the accession ‘Pourpre Hollande’, three accessions from Eastern Europe (‘Plovdivski’ from Bulgaria, ‘A 117-15’ from Hungary, ‘VL25B’ from Romania), the Israeli accession ‘Kfar Hanania’, and ‘UK 215AG12’ from Central Asia.

**Table 1.**
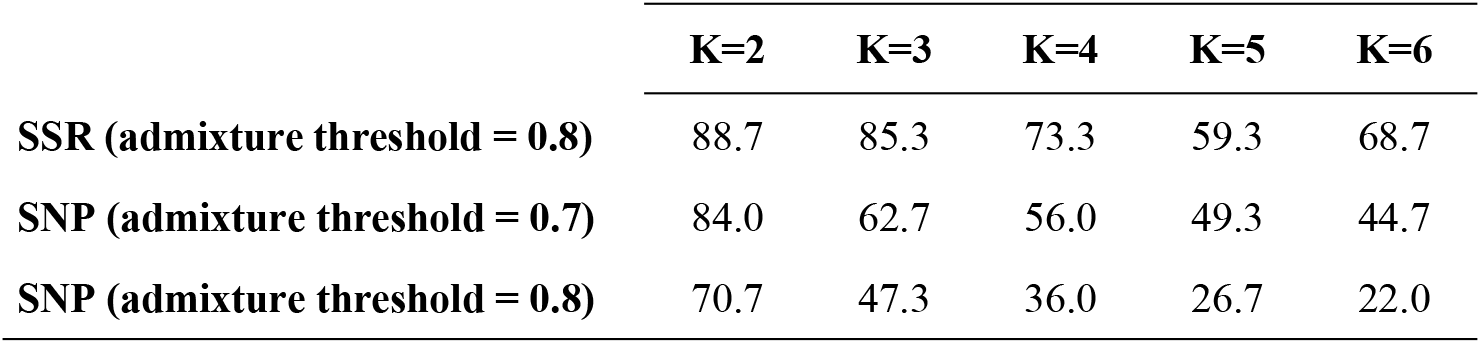
Percentage of population assignment from K=2 to K=6, using SSR and SNP markers.

Using SNP markers and a threshold of 0.7, we found 24 admixed accessions (84% of assignment), including 9 of the 17 found using SSR markers. From these accessions, 7 (‘A 117-15’, ‘Fernette’, ‘Pourpre Hollande’, ‘Amigo’, ‘Chico’, ‘Forde’, ‘Gillet’) are now clustered into the group B, and ‘UK 215AG12’ is now in the group A. In addition, 14 accessions from group A, and 1 from group B, based on SSR clustering, are found admixed using SNP markers (Figure 3). So, only 23/150 accessions (15%) are differently clustered. In any case, we did not find any group exchange from A to B, or from B to A, by comparing the clustering based on SSR or on SNP markers.

**Fig 3.**
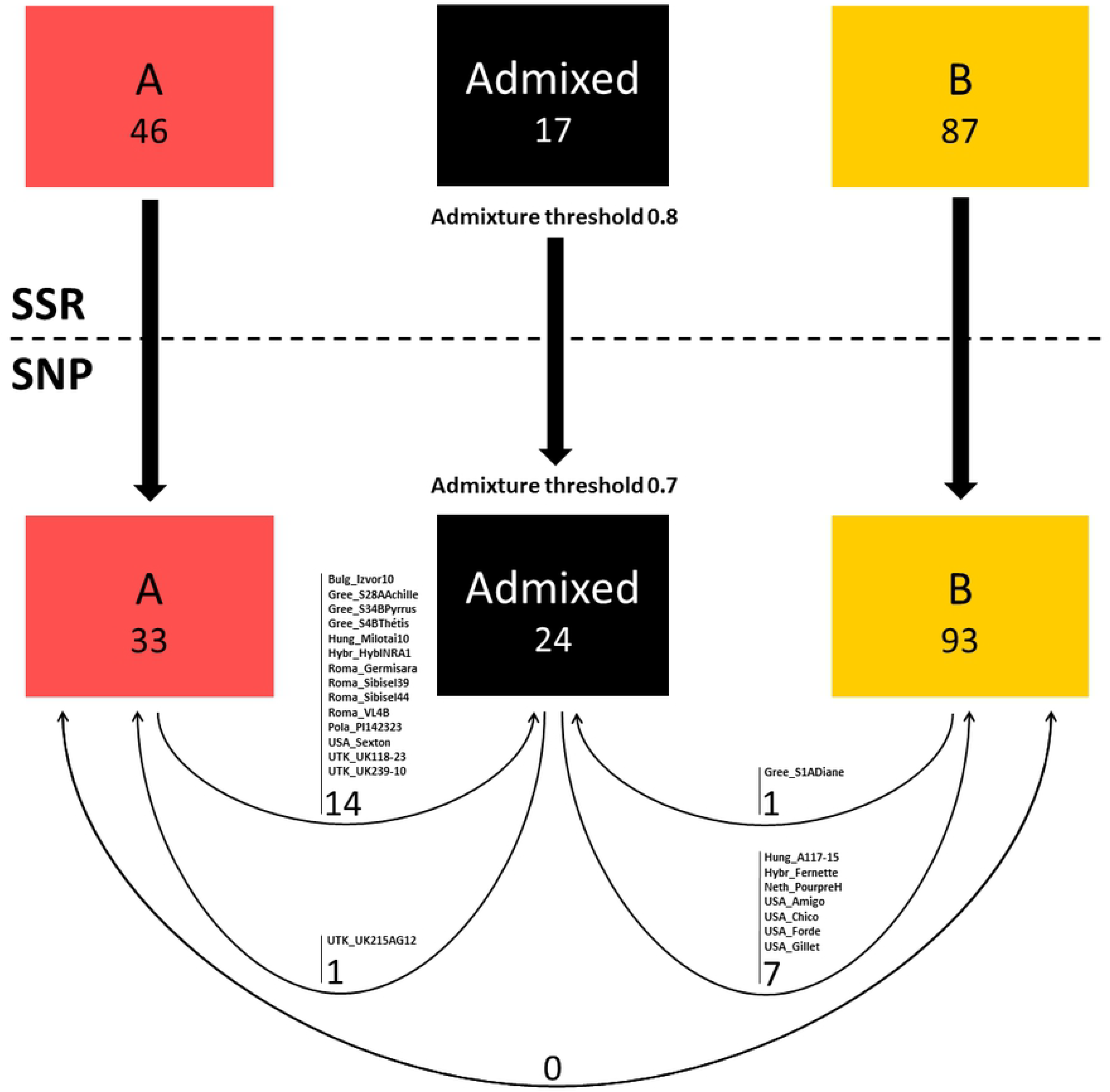
SNP-based clustering results, compared to SSR-based clustering results. For a threshold of 0.7 for admixture, SNP markers clustered 14 accessions found in the group A, and one found in the group B, compared to SSR markers, into the admixed group. Conversely, SNP markers clustered one and seven accessions found admixed, compared to SSR markers, into the groups A and B, respectively. There is no clustering exchange between the groups A and B, comparing both methods.

When using a threshold of 0.8 for the SNPs, the percentage of population assignment for K=2 is 70.7%, and drops to 47.3% for K=3, whereas it is still high for a threshold of 0.7 (62.7%) (Table 1). In addition, we ran a Spearman rank correlation test for K=2 and we found that the clustering between SSR and SNP markers is highly correlated, up to 84%.

### Comparison of the first level of structure with PCoA results

The PCoA constructed in 2D and 3D show the clustering of the 150 accessions following results obtained from K=2 (Figure 4). For K=2, the PCoA results are in agreement the structure based on SSR or SNP markers, since the scatterplots for the groups A and B are well defined by the first principal component. The admixed accessions are positioned mainly between the groups A and B. Moreover, for both methods, the three Manregian walnuts (‘Chase C7’, ‘Wepster W2’ and ‘Adams 10’), which are trees originated from seed collected in northeastern China, are isolated and found to be genetically diverse. Regarding the percentage of explained variation, they are also comparable using both types of markers. The first three axes (x, y, and z) explain 21.86% of the cumulative variation for SSRs, and 14.91% for SNPs.

**Fig 4.**
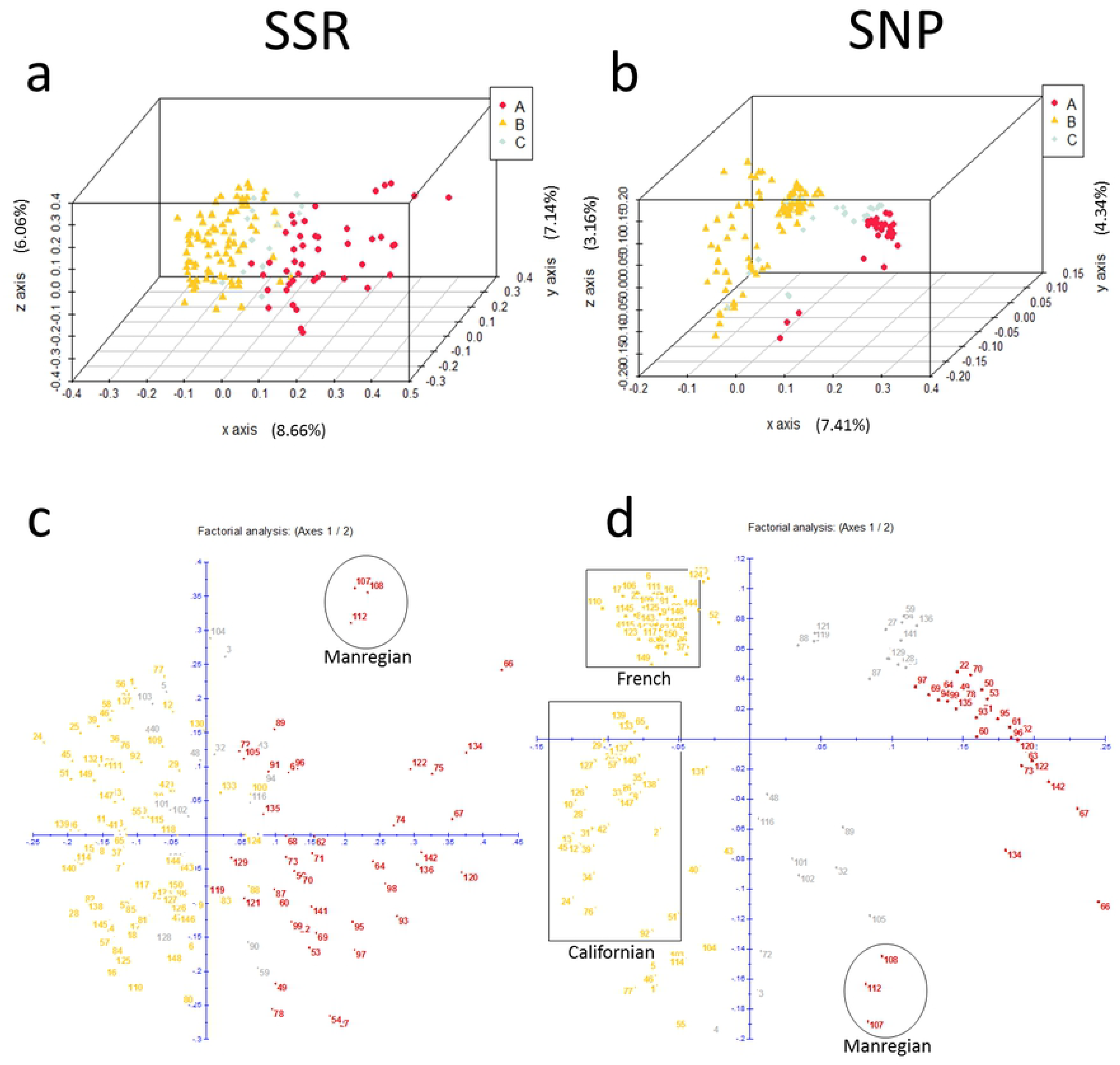
Principal Coordinate Analysis scatterplots. PCoA were constructed in 3D using a) SSR, and b) SNP markers, and in 2D using c) SSR, and d) SNP markers. The 150 accessions are colored following K=2 results: group A in red, group B in yellow, and admixed in grey.

However, we found some differences between the two types of marker. Using SSR markers, the scatterplot corresponding to the group A is more extensive, whereas the group B is more scattered using SNP markers, and besides, this group is split in two, showing the French accessions on one side (‘Candelou’, ‘Grandjean’, ‘Lalande’, ‘Quenouille’, ‘Lub’, ‘Chaberte’, etc.) and the Californian on the other (‘Carmelo’, ‘Howe’, ‘Trinta’, ‘Tehama’, ‘Waterloo’, ‘Hartley’, etc.), using the second principal component.

### Comparison of the first level of structure with grouping trees results

The Neighbor-joining method implemented in “DARwin 6.0.14” permitted to construct grouping trees with the 150 accessions (Figure 5). The main branching groups of the trees obtained with both markers, are in agreement with the structure results (K=2), since they are mainly defined by the groups A and B. The two accessions ‘Jin Long 1’ from China and ‘PI 15 95 68’ from Afghanistan have a long length branch, indicating a high level of genetic diversity, for both methods. However, few differences were detected between the structure and the tree using SSR markers: 9 accessions of the group A are found in the branching group mainly corresponding to the group B (‘IR 13-1’, ‘Hybrid INRA 3’, ‘UK 41-17’, ‘S 4 B Thétis’, ‘Sexton’, ‘Milotai’, ‘PI 14 23 23’, ‘Z 53’, ‘PI 15 95 68’). Using SNP markers, this is the case for three accessions (‘PI 15 95 68’, ‘Wepster W2’, ‘Adams 10’).

**Fig 5.**
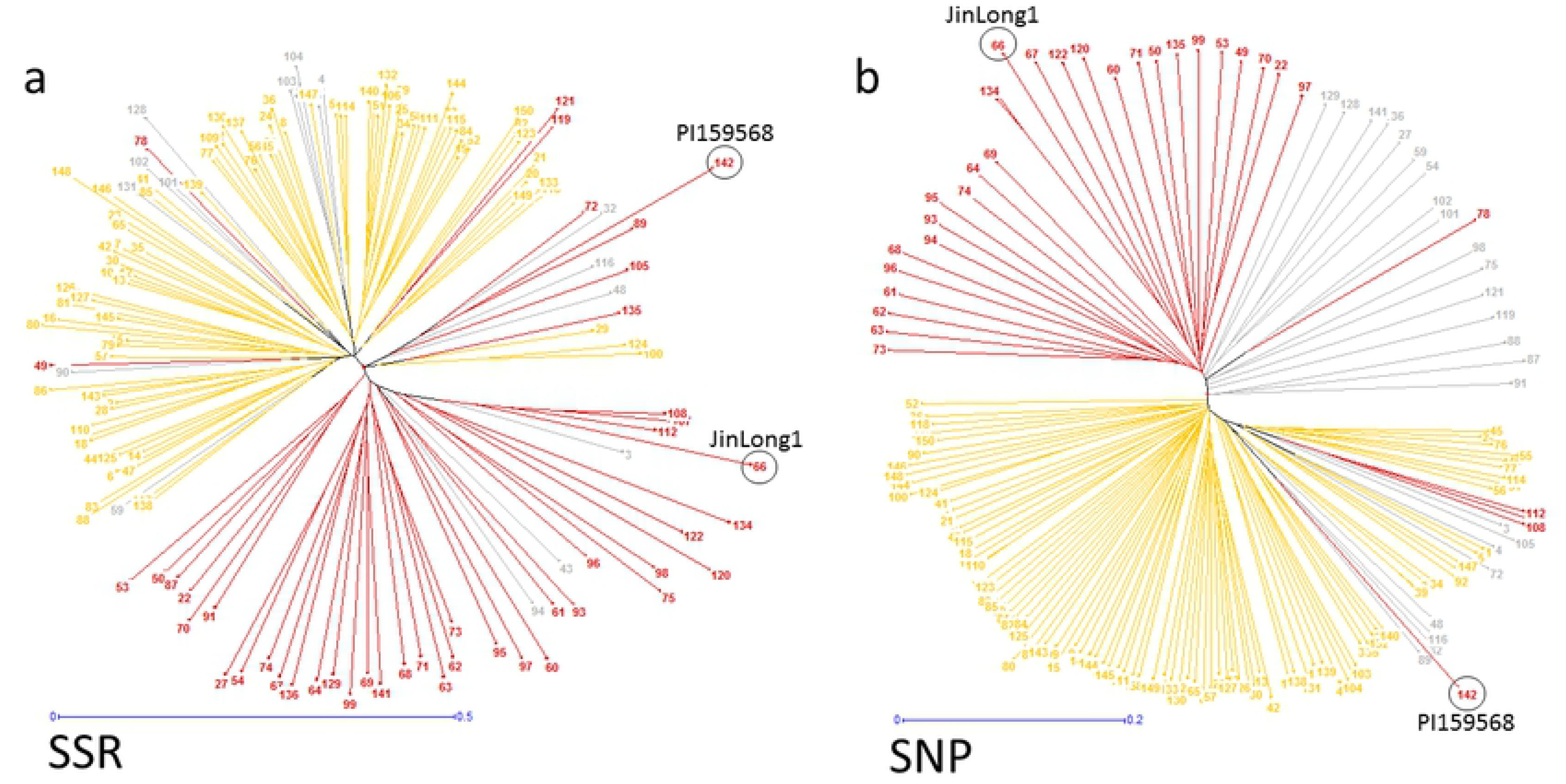
Neighbor-joining trees. Trees were constructed using a) SSR, and b) SNP markers. The 150 accessions are colored following K=2 results: group A in red, group B in yellow, and admixed in grey.

### Structure results from K=3 to K=6

We then inferred the individual admixture coefficients of the 150 accessions from K=3 to K=6 for both methods (Table S2). Interestingly, results are slightly contrasted depending on the markers. For K=3 (group C), the SSR markers highlight for example the French and Californian modern hybrids (Figure S1). Using the SNP markers, the group C contains the French modern hybrids but also the French landraces only coming from South-Est such as ‘Franquette’, ‘Mayette’, ‘Meylannaise’, ‘Romaine’, and ‘Parisienne’. It contains also all the Californian accessions, not only the modern hybrids. For K=4 (group D), the SSR markers mainly emphasize the Californian accessions with ‘Payne’ within their pedigree (‘Ashley’, ‘Chico’, ‘Chandler’, ‘Howard’, ‘Marchetti’), and the French landraces only coming from South-West such as ‘Grosvert’, ‘Ronde de Montignac’, ‘Lalande’, and ‘Solèze. We also found the French modern hybrids (Figure S2). Results are very similar using the SNP markers since the group D contains also ‘Payne’ pedigree’s accessions and the French hybrids. For K=5 (group E), the SSR markers highlight ‘Lu Guang’ from China, ‘Sopore’ from India, the accessions from Romania, and the accessions from Central Asia (Uzbekistan, Tajikistan, Kyrgyzstan). The group D is now for the accessions with ‘PI 15 95 68’ from Afghanistan in their pedigree such as ‘Serr’ and ‘Tulare’ (Figure S3). Using SNP markers, the group E contains the Japanese and Chinese accessions except ‘Lu Guang’, the French modern hybrids, ‘Gillet’ and ‘Sexton’, two Californian modern hybrids with Chinese pedigree, and ‘Lara’. For K=6 (group F in black), the SSR markers emphasize ‘PI 15 95 68’ pedigree’s accessions, as it was the case for K=5 with the group D. The group F also contains ‘Sexton’ with Chinese pedigree, ‘Kfar Hanania’ from Israel, and ‘S 4 B Thétis’ from Greece (Figure S4). The group D now has the French landraces from South-West and ‘Payne’ pedigree’s accessions. Using SNP markers, the group F contains the Chinese and Japanese accessions, ‘EAA 6’ from Greece, and the accessions from Central Asia.

### Core collection construction using SSR and SNP markers

We constructed core collections of 50 accessions with both methods, using SSR and SNP markers. With the method based on the “maximum length sub tree” function of “DARwin 6.0.14”, 32/50 accessions are in common between the data sets based on SSR or SNP markers (Table 2). The accessions belong mainly to the group A, from Eastern Europe and Asia, known to be more diverse (29/50 using SSRs, 20/50 using SNPs). They both include the Iranian accessions, the Indian ‘Sopore’, the Bulgarian ‘Izvor 10’ and ‘Plovdivski’, and several accessions from the Botanical Garden of Kiev. Regarding the French accessions, both markers kept the particular accessions ‘RA 1195’, a weeping tree, and ‘RA 1100’, a tree particularly resistant to frost. Only ‘Corne’ and ‘Marbot’ were kept as French landraces using SSRs, and ‘Chaberte’ using SNPs. Similar results were observed with the “entry-to-nearest entry” method with also 32/50 accessions in common between the data sets based on SSR or SNP markers.

**Table 2.**
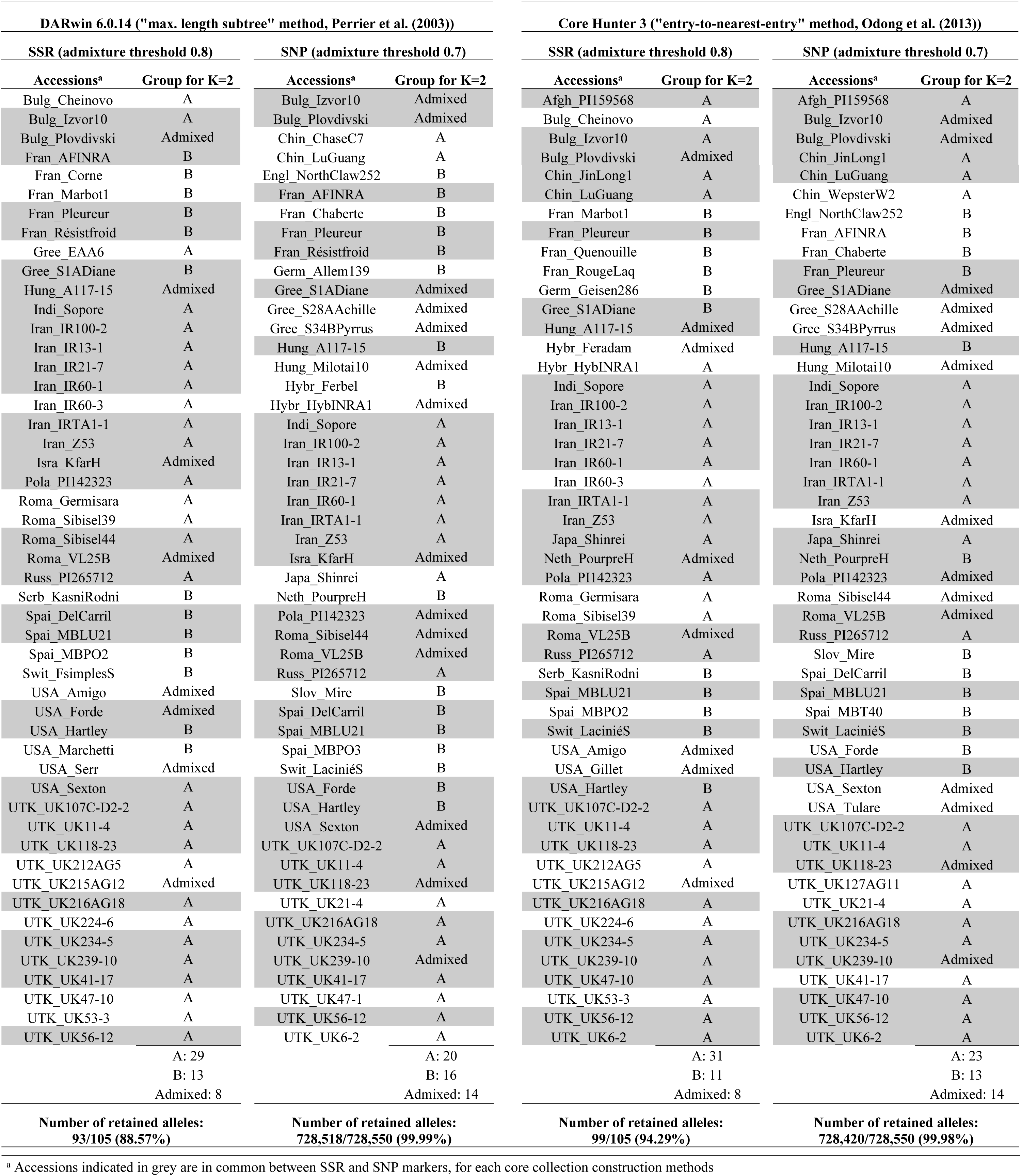
Construction of the core collections (n=50) using SSR and SNP markers, and two different methods of construction.

Moreover, the consistency of the results between core collection construction methods was checked for both markers. Using SSR markers, 37/50 accessions are in common between the two methods, and 43/50 accessions using SNP markers. We estimated the number of alleles retained in each core collection. For SSR markers, we retained 88.57% and 94.29% of the 105 total alleles found within the entire collection, using the “maximum length subtree” and the “entry-to-nearest entry” methods, respectively. Using SNP markers, we retained 99.99% and 99.98% of the 728,550 total alleles found, with the same methods, respectively (Table 2).

### Comparison of three sets of SNPs for PCoA assessment

In addition, for PCoA assessment, we compared the entire set of 364,275 SNPs with a subset of SNPs filtered for LD, with a threshold of r²=0.2 (Figure 6a). We then retained 24,422 SNPs, or 6.7% of the entire set. Interestingly, the results found for both datasets are strongly similar, with the scatterplots well distinguished, according to the K=2 results (Figure 6b). We still distinguish the French accessions from the Californian accessions within the group B. The main difference is that the variance is better explained by the first three axes using the LD-pruned set (18.90% vs. 14.91%). By comparing the entire set with a random subset of 100 SNPs, a range comparable to the total number of SSR alleles, we found that the scatterplots are less well defined, but still in agreement with K=2 results (Figure 6c). In this case, we cannot distinguish the French accessions from the Californian accessions within the group B.

**Fig 6.**
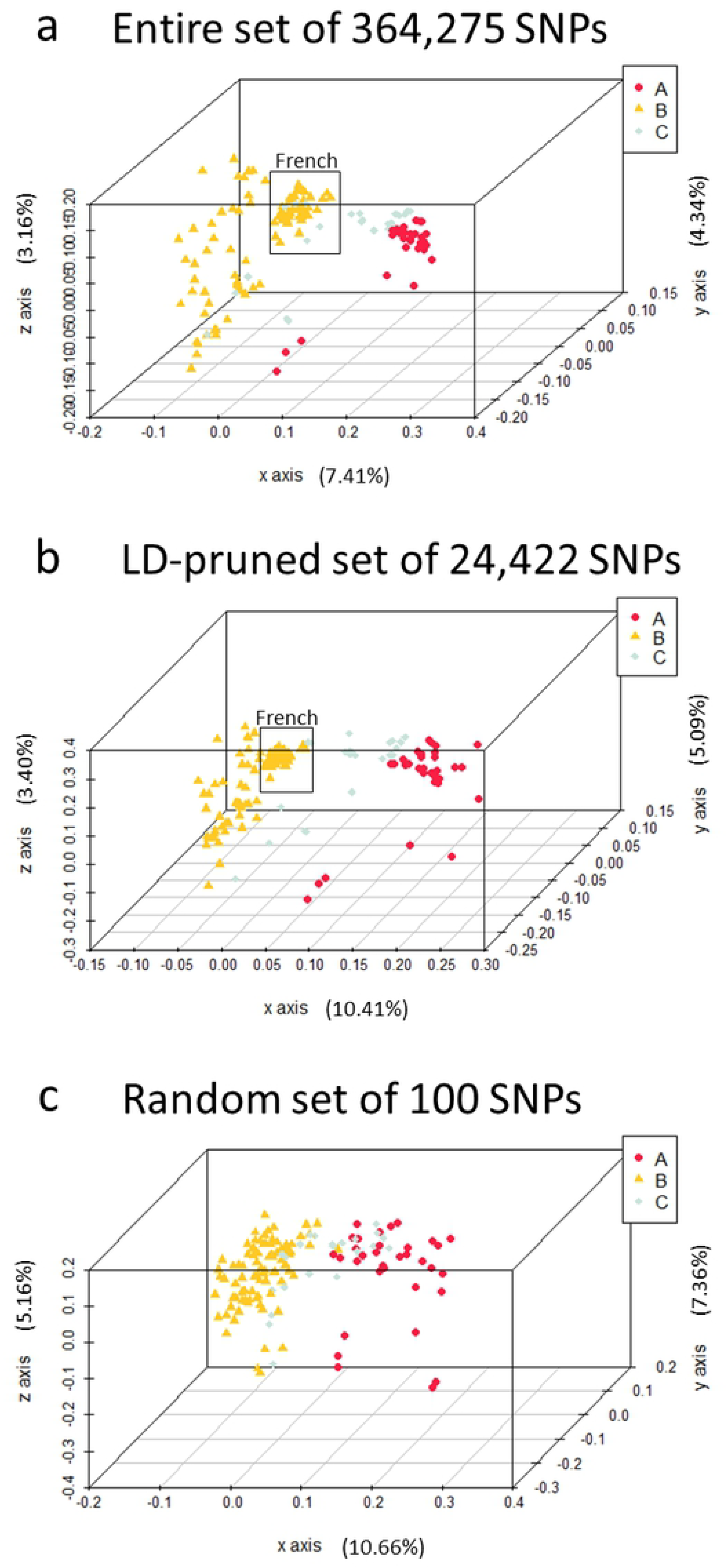
Comparison of SNPs set for Principal Coordinate Analysis. PCoA were constructed using a) the entire set of 364,275 SNPs, b) the LD-pruned subset of 24,422 SNPs, and c) the random set of 100 SNPs.

## Discussion

### The use of either SSR or SNP markers shows comparable results for structure analyses

By genotyping the panel of 150 *J. regia* accessions using 13 SSRs and more than 300,000 SNPs, we obtain similar and comparable results. Both types of marker showed a first level of structure for K=2, with no exchange of accession between the main groups (A and B), which are related to the geographical origin of the accessions. The exchanges only concern accessions that switched from one group to the admixed cluster, or vice versa. From K=3 to K=6, the results remain highly comparable with the highlighting of substructures linked, for instance, to ‘Payne’ pedigree for the Californian modern hybrids, or to geographical area (South-West vs. South-Est) for French landraces. Then, when compared structure results for K=2 with PCoAs, and with grouping trees, results are still consistent. More precisely, we found as the most diverse accessions, using both markers, the three Manregian in PCoA, and ‘PI 15 95 68’ and ‘Jin Long 1’ using trees, all coming from Asia. When considering the LD-pruned set of 6.7% of the entire set of SNPs, PCoA showed consistent clustering patterns and also with those using the 13 SSRs.

This kind of findings was previously observed in other works, except that the number of needed markers would be different to obtain a comparable resolution power. For example, the broad patterns of PCoA were similar using 36 SSRs with 2.2 alleles per locus in average and 36 SNPs in 375 Indian accessions of rice [13]. But, in local accessions of cowpea from East African countries, similar clustering patterns were found using more SNPs than SSRs (151 vs. 13) [19]. Also in jujube, within a core-collection of 150 accessions, only 18 (12%) were classified into different groups based on the results of structure analysis using 24 SSRs and 4,680 SNPs [21]. Within various inbred maize lines, SSRs performed better at clustering accessions into groups using about 10 times more SNPs [14,16]. Other works suggest in the same vein the use of three times [18], or seven to 11 times more SNPs to obtain comparable informative results [15]. In our study, we have taken care to choose highly polymorph and robust SSRs developed for *J. regia*, by reviewing the literature [37–39]. When working with biallelic markers such as SNPs, it is known that the genetic distances can be equivalent to those calculated with SSRs, using this formula: *n*(*k*-1), where *k* is the average number of alleles per locus, and *n* is the number of loci [40]. With 13 loci and 8.1 alleles per locus in average (four times of that for SNPs), we need in theory [13*(8.1-1)] = 93 SNPs to obtain equivalent genetic distances in our panel. Our findings based on PCoA clustering using the random set of 100 SNPs are consistent with this number of SNPs.

Moreover, SNPs globally tend to give higher proportions of inferred admixture, as observed in sunflower [17]. Regarding the admixture thresholds chosen, the results found when compared 0.7 or 0.8 for SNPs also show that 0.7 is clearly more suitable to obtain comparable structure. In this respect, we found that the percentage of assignment decreases as the K increases, particularly for the SNPs. Such differences were also reported in maize [16] and grape [20].

### The choice of marker type will depend on the needed task related to germplasm conservation or utilization

The management of PGR comprises the following main steps: their conservation, which consists in acquisition of plant material (by the collection or the protection of reserves *in situ*, or by exchange of *ex situ* material), their maintenance (such as storing and propagation), their characterization (based on both genotype and phenotype) and their utilization for research, breeding programs or production [41]. Due to the increasing availability of genomics tools, we talk about “genoplasmics”, a cross-disciplinary field which aims to use genomics in germplasm management [42]. But undoubtedly, the choice of SSR or SNP markers will depend on the purposes.

For obvious reasons, a first choice criterion could be the cost of genotyping. New genomics technologies have a cost decreasing dramatically for past years. Apart from the DNA extraction, the cost of SNP vs SSR genotyping was about 2,600 times less in our study. For guidance only, we paid 8.8 USD$ for 13 SSR loci per sample (0.7 USD$/locus/sample) and 98.9 USD$ for one array of 364,275 robust loci per sample (2.7E-4 USD$/locus/sample). However, the SSR genotyping is often more “flexible”, since we could choose precise numbers of loci and samples. In the case of a SNP array, all the loci available are assayed, for a 96-well DNA plate. In addition, only 59.8% of the available SNPs on the array were usable for the analyses after quality control.

A second choice criterion could be the nature of the plant material managed, particularly if the researcher works only on one crop, or also with its wild species. In our case, the SSRs used were also highly transferable into wild species of the genus *Juglans* spp. [24]. On the contrary, the Axiom™ *J. regia* 700K SNP array used is valid on the cultivated species *J. regia* only and failed on our few wild species accessions tested. But this case is not a general case. In [20], the set of 384 SNPs used permitted to analyze 2,273 accessions of grape (*Vitis* spp.), including cultivated grapevines (*V. vinifera* ssp. *sativa*), wild grapevines (*V. vinifera* ssp. *sylvestris*) and non-*vinifera Vitis* species used as rootstocks.

A third reason to consider either marker could be the main goal of the genotyping. Well-chosen neutral SSRs would be sufficient for “simple” population structure and relationships determination, particularly in the first steps of germplasm management, since the computational time for analyzes is lower. But SNPs, likely to be associated with functional variation, would be preferred for genome-wide association study purpose, and may have higher resolution for relatedness estimations. To construct core-collections, our results were similar, knowing that it is easier to keep the entire allelic diversity using SNPs.

### The preservation of the allelic diversity must be compatible with the preservation of the phenotypic variability

To our knowledge, the literature is missing concerning the marker type that should be ideally used to construct an effective core-collection. In light of our results, we found 64% of similarity between the two marker types, for the chosen accessions. They both preferentially kept accessions from East Europe and Asia, as expected, because of their global higher genetic diversity, and both markers helped to understand that French landraces have a moderate level of genetic diversity. Only the French landraces ‘Corne’, ‘Marbot’ and ‘Chaberte’ were kept using SSRs or SNPs. Knowing that French landraces represent 20% of the entire plant material panel, these findings confirm that their diversity is moderate. Moreover, the four core-collections constructed kept at least 88.57% of the allelic diversity.

In parallel with the preservation of the allelic diversity, it is also necessary to take into account the phenotypic variability. The INRAE walnut germplasm collection contains some accessions with unusual traits, such as weeping branches, laciniate leaves or purple foliage, which may be used for ornamental purposes. Interestingly, the accession with weeping branches ‘RA 1195’ is in the four core-collections, and the accessions with laciniate leaves or purple foliage are in three of them. Based on chronological phenotypic data available [43] and new data acquired, we also looked if the core-collections contain a high or small range of the variability of some important traits. Let us take the example of a trait related to the phenology, crucial for climate change adaptation, the budbreak date. In average, the ten earlier accessions are ‘Early Ehrhardt’, ‘Mire’, ‘Payne’, ‘Serr’, ‘Kfar Hanania’, ‘IR 60-1’, ‘Sopore’, ‘Z 53’, ‘Ashley’ and ‘Lu Guang’, with a range of budbreak date from 65 to 75 Julian days. Five or six of them are found depending on the core-collection. On the contrary, none of the ten later accessions (‘Fertignac’, ‘Le Bordelais’, ‘St Jean n°1’, ‘Lalande’, ‘Candelou’, ‘Maribor’, ‘Semence Comité Dordogne’, ‘Ronde de Montignac’, ‘Culplat’ and ‘Romaine’), with a range of budbreak date from 110 to 122 Julian days, is found in the four core-collections. Here is the limit of a management only based on molecular data: the genetic diversity kept will not necessary keep the phenotypic variability.

## Conclusion

In our comparison using 150 *J. regia* accessions, both SSR and SNP markers were globally equally appropriate, for both structure analysis and core-collection construction. It is therefore important to consider the task of the germplasm management to choose the most appropriate marker. In general, SSR markers are suitable to obtain a global idea of the structure of a germplasm. However, if the goal is to use the collection for genomics analysis such as genome-wide association studies (GWAS), a high number of SNPs are required. Typically, that is what we did for the management of the INRAE walnut germplasm collection. From the 217 *J. regia* accessions available in the collection, we choose a set of 170 accessions, based on SSR markers, to perform then GWAS using the Axiom™ *J. regia* 700K SNP array. Those SSR markers have permitted to set aside synonym and/or redundant accessions.

## Acknowledgments

We want to thank the Fruit Tree Experimental Unit of the INRAE in Toulenne and the *Prunus/Juglans* Genetic Resources Center for the maintenance of the collection and for helping us to collect the samples. We acknowledge the BioGEVES laboratory for DNA extraction and SSR genotyping, and ThermoFisher for SNP genotyping. Then, the CTIFL, holder of the project “INNOV’noyer”, in partnership with the INRAE of Bordeaux, want to thank the “Région Nouvelle-Aquitaine”, and “Cifre” convention of “ANRT” (Agence Nationale de la Recherche et de la Technologie). It is also important to note that the project is supported by the “Agri Sud-Ouest Innovation” competitiveness cluster. Finally, we would like to thank the late Eric Germain, former head of the breeding program at the INRAE of Bordeaux from 1977 to 2007. His remarkable work, then continued by Francis Delort, has given us the opportunity to study a rich set of plant material.

## Supporting information

**S1 Fig. The bar plots showing the individual admixture coefficients of the 150 accessions for K=3.** Structure was assessed a) using SSR, and b) using SNP markers.

**S2 Fig. The bar plots showing the individual admixture coefficients of the 150 accessions for K=4.** Structure was assessed a) using SSR, and b) using SNP markers.

**S3 Fig. The bar plots showing the individual admixture coefficients of the 150 accessions for K=5.** Structure was assessed a) using SSR, and b) using SNP markers.

**S4 Fig. The bar plots showing the individual admixture coefficients of the 150 accessions for K=6.** Structure was assessed a) using SSR, and b) using SNP markers.

**S1 Table. List of the 150 accessions from INRAE walnut germplasm collection.**

**S2 Table. Q-matrices showing the individual admixture coefficients of the 150 accessions from K=2 to K=6, using SSR and SNP markers.**

**S3 Table. SSR genotyping data set**

The SNP genotyping data set in “hapmap” format is freely and openly accessed on the “Portail Data INRA” repository, via the identifier “INRA’s Walnut Hapmap” and the following Digital Object Identifier (DOI): https://doi.org/10.15454/XPKII8. The dataset called “GWAS_hapmap.txt” is related to a GWAS panel of 170 accessions, including the 150 accessions of this study. The additional file called “List of ID.tab” allows to link the array identifier name of the accessions with the identifier name used in this study.

